# Population-wide Screening for Germline Variants of Hereditary Cancer Genes in 12K Unselected Japanese Colorectal Cancers and 27K Controls

**DOI:** 10.1101/2020.03.15.989947

**Authors:** Masashi Fujita, Xiaoxi Liu, Yusuke Iwasaki, Chikashi Terao, Sadaaki Takata, Chihiro Inai, Tomomi Aoi, Kazuhiro Maejima, Makoto Hirata, Yoshinori Murakami, Yoichiro Kamatani, Michiaki Kubo, Kiwamu Akagi, Koichi Matsuda, Hidewaki Nakagawa, Yukihide Momozawa

## Abstract

**Background & Aims:** Colorectal cancer (CRC) is one of the most common cancers in Western countries and Japan. Currently, a few % of CRCs can be attributed to recognizable hereditary germline variants of known CRC susceptibility genes, predominantly the DNA mismatch repair genes. To establish a universal screening strategy for hereditary CRCs, it is necessary to explore the prevalence of hereditary CRC and pathogenic variants of multiple cancer-predisposing genes in non-European populations.

**Methods:** We analyzed the coding regions of 27 cancer-predisposing genes, including mismatch repair genes, *APC*, and *BRCA1/2*, in 12,503 unselected Japanese CRC patients and 23,705 controls aged ≥ 60 years without any personal or family history of cancer by target sequencing and genome-wide SNP chip data. Their clinical significance was assessed using ClinVar and the guidelines by the American College of Medical Genetics and Genomics and the Association for Molecular Pathology (ACMG/AMP).

**Results:** We identified 4,804 variants in the 27 genes and annotated them as 397 pathogenic variants, 941 benign variants, and 3,466 variants of uncertain significance, of which 43.6% were registered in neither ClinVar nor dbSNP. In total, 3.3% of the unselected CRC patients and 1.5% of the controls had a pathogenic variant of the 27 genes. The pathogenic variants of *MSH2* (odds ratio (OR) =18.1), *MLH1* (OR=8.6), *MSH6* (OR=4.9), *APC* (OR=49.4), *BRIP1* (OR=3.6), *BRCA1* (OR=2.6), *BRCA2* (OR=1.9), and *TP53* (OR=1.7) were significantly associated with CRC development in the Japanese population (*P*-values < 0.01, FDR<0.05). Furthermore, we confirmed copy number variants (CNVs) of *MSH2/EPCAM, MLH1*, and *APC* by multiplex ligation-dependent probe amplification (MLPA) and quantitative PCR in this cohort (n = 23), including whole gene duplications of *MSH2* and *APC.* These pathogenic variants were significantly associated with the diagnostic age and personal/family history of other types of cancer. In total, at least 3.5% of the Japanese CRC population had a pathogenic variant or CNV of the 27 cancer-predisposing genes.

**Conclusions:** This is the largest study of CRC heredity in the Asian population and would contribute to the development of guidelines for genetic testing and variant interpretation for heritable CRCs. Universal screening for CRC risk should be assessed in multiple genes, including *BRCA1/2* and *BRIP1*. These data would facilitate risk assessment of cancer and optimize the screening strategy.

## INTRODUCTION

Colorectal cancer (CRC) is one of the most common cancers in Western countries and Japan, and approximately 20% of CRC cases have affected relatives of CRCs or other types of cancers, suggesting relatively higher hereditability (1-3). Common risk loci explain up to 8% of CRC heritability. More than 50 susceptibility variants have been identified by genome-wide association studies (4, 5), but they explain only 1–4% of the underlying genetic variation. Familial or hereditary CRCs have been one of the most common subjects by genomic analysis, and a few percent of CRCs can be attributed to recognizable hereditable rare germline variants of cancer susceptibility genes (1, 2). Lynch syndrome, caused by germline mutations in the DNA mismatch repair (MMR) genes *MLH1, MSH2, MSH6*, and *PMS2* (1, 2, 6, 7), is the most common cause of hereditary CRC and accounts for 1–3% of patients with CRC. Patients with tumors exhibiting characteristics of MMR deficiency are more likely to have Lynch syndrome; therefore, professional guidelines recommend that all patients with CRC undergo tumor screening for Lynch syndrome, with referral to genetic counseling for those with MMR deficiency (6, 7). The National Comprehensive Cancer Network (NCCN) Guidelines for Genetic/Familial High-Risk Assessment also suggest that all CRC patients younger than 50 years old should consider genetic testing for Lynch syndrome. Another main genetic player in hereditary CRC is *APC* (1, 2). Germline pathogenic or truncated variants of *APC* are responsible for familial adenomatous polyposis (FAP), and somatic mutation of *APC* is responsible for most sporadic CRC or precursor adenoma polyps. *APC* is a gatekeeper for CRC development in multistep carcinogenesis (8). However, the FAP phenotype is quite rare and distinct at the population level, and screening for germline variants of the *APC* gene is not recommended for common CRCs. The prevalence of other hereditary cancer syndromes among patients with CRC is largely unknown because previous studies on other genes are limited and have been confined to selected (high-risk) patient populations.

With advances in next-generation sequencing (NGS), genetic testing for hereditary cancers has shifted from phenotype-specific single gene assessment to broad panels providing simultaneous assessment of multiple genes implicated in various hereditary cancer syndromes (10, 11). Multigene panel testing for hereditary CRC is feasible, timely, and more cost-effective than single-gene testing at present. However, these genetic tests identify a large number of variants, most of which are clinically variants of uncertain significance (VUS) and sometimes ethnicity-specific variants, complicating the interpretation of the results. The prevalence of hereditary cancers such as Lynch syndrome varies significantly in different populations, suggesting that ethnic diversity might play an important role in hereditary diseases, and variant data of the Asian population is still limited (12). Screening for Lynch syndrome and other hereditary cancers has been mainly performed in Western populations, and data regarding other ethnicity patients are scarce, leading to difficulties in interpreting variants of cancer-predisposing genes in Japan and the Asian population. Hence, it is necessary to have a large-scale variant dataset for a specific ethnic group.

In this study, to explore the prevalence of hereditary CRC and pathogenic variants of cancer-predisposing genes, we sequenced almost all coding regions of 27 cancer-predisposing genes in 12,503 unselected Japanese CRC patients and 23,705 controls in the biobank. We then assessed these variants following the American College of Medical Genetics and Genomics (ACMG) guidelines and demonstrated the prevalence of hereditary CRC and variants of cancer-predisposing genes in the Japanese population. These findings facilitate risk assessment of CRCs, direct clinical management, and optimize the screening strategy for CRC and other types of cancer in Japanese and other populations.

## Methods

### Study population

We obtained all study samples from BioBank Japan (13, 14), which is a multi-institutional, hospital-based registry that collects blood DNAs and clinical information from patients with various common diseases, including CRC from all over Japan between 2003 and 2018. Clinical characteristics of CRC cases and controls were collected by interview or medical record survey using a standard questionnaire at the point of entry to Biobank Japan (15). We selected 12,606 CRC cases from this cohort and used the same 23,780 controls aged ≥ 60 years with no personal or family history of cancer from our previous study on breast cancer [16]. All participants provided written informed consent. The study was approved by the ethics committees of the Institute of Medical Sciences, The University of Tokyo, and the RIKEN Center for Integrative Medical Sciences.

### Sequencing and bioinformatic analysis

We selected 27 genes, including the 12 genes recommended by the NCCN guidelines whose rare germline variants were reported to show high penetrance for CRC and hereditary cancers (17). We analyzed the complete coding regions and 2-bp flanking intronic sequences of all 27 genes, except exons 10–15 of *PMS2*, (84,822 bp) by a multiplex PCR-based target sequence method (16, 18). We called single nucleotide variants (SNVs) and insertion or deletion (INDELs) of each individual separately using UnifiedGenotyper and HaplotypeCaller of GATK, as described previously (16, 18). Genotypes for all individuals were jointly determined for each variant based on the sequencing read ratio of reference and alternative alleles. We assigned homozygotes of the reference allele, heterozygote, or homozygote of the alternative allele, when the alternative allele frequency fell in the range of 0-0.15, 0.25-0.75, or 0.85-1, respectively. Finally, we identified 4,804 genetic variants in 12,503 patients and 23,705 controls, and 99.952% of the target region was covered by ≥ 20 sequence reads.

### Annotation of variants

We assigned clinical significance (pathogenic, benign, or uncertain) for all variants, as in our previous study (16, 18) with some modifications. Briefly, we determined clinical significance using the American College of Medical Genetics and Genomics and the Association for Molecular Pathology (ACMG/AMP) guidelines (19) as well as the pathogenicity assertions registered in ClinVar. We considered variants as pathogenic based on classification as pathogenic by the ACMG/AMP guidelines and/or classification as pathogenic in ClinVar (criteria for classifying variants is **Supplementary Table 1 and 2**). Variants not registered in ClinVar on 20^th^ May 2019 were considered novel.

**Table 1.** Demographic and clinical characteristics of study participants.

|  | CRC patients, No. (%) | Controls, No. (%) |
| --- | --- | --- |
| Number of subjects | 12,503 | 23,705 |
| Mean Age at entry, y (SD) | 67.6 (10.3) | 71.1 (7.3) |
| Mean age at CRC diagnosis, y (SD) | 65.1 (10.5) | -- |
| Sex |  |  |
| Male | 7,871 (63.0) | 12,477 (52.6) |
| Female | 4,632 (37.0) | 11,228 (47.4) |
| Personal history of gastric cancer | 295 (2.4) | 0† (0.0) |
| Personal history of prostate cancer | 132 (1.1) | 0† (0.0) |
| Personal history of breast cancer | 138 (1.1) | 0† (0.0) |
| Personal history of endometrial cancer | 46 (0.4) | 0† (0.0) |
| Personal history of ovarian cancer | 41 (0.3) | 0† (0.0) |
| Personal history of FAP | 2 (0.0) | 0† (0.0) |
| Personal history data missing | 2 (0.0) | 0† (0.0) |
| Family history of CRC | 1,889 (15.1) | 0† (0.0) |
| Family history of gastric cancer | 2,628 (21.0) | 0† (0.0) |
| Family history of prostate cancer | 300 (2.4) | 0† (0.0) |
| Family history of breast cancer | 660 (5.3) | 0† (0.0) |
| Family history of endometrial cancer | 122 (1.0) | 0† (0.0) |
| Family history of ovarian cancer | 69 (0.6) | 0† (0.0) |
| Family history of FAP | 3 (0.0) | 0† (0.0) |
| Family history data missing | 2 (0.0) | 0† (0.0) |
| Anatomic site of CRC |  |  |
| Right-side colon | 3,471 (29.8) | -- |
| Left-side colon and rectum | 8,025 (68.9) | -- |
| Both sides of colon | 158 (1.4) | -- |
| Multiple CRC lesions |  |  |
| Yes | 757 (7.5) | -- |
| No | 9,290 (92.5) | -- |
| Lymph node metastasis |  |  |
| Yes | 2,582 (43.0) | -- |
| No | 3,426 (57.0) | -- |
| Distant metastasis |  |  |
| Yes | 586 (12.3) | -- |
| No | 4,186 (87.7) | -- |
| Histology |  |  |
| Adenocarcinoma |  |  |
| Papillary adenocarcinoma | 64 (0.6) | -- |
| Tubular adenocarcinoma | 9,672 (89.4) | -- |
| Poorly differentiated adenocarcinoma | 315 (2.9) | -- |
| Mucinous adenocarcinoma | 162 (1.5) | -- |
| Signet-ring cell carcinoma | 16 (0.1) | -- |
| Adenocarcinoma, NOS | 395 (3.6) | -- |
| Adenosquamous carcinoma | 19 (0.2) | -- |
| Squamous cell carcinoma | 23 (0.2) | -- |
| Carcinoid tumor | 43 (0.4) | -- |
| Others | 113 (1.0) | -- |

†Controls with no personal history or family history of cancers were selected for this study. NOS, not otherwise specified.

**Table 2.**
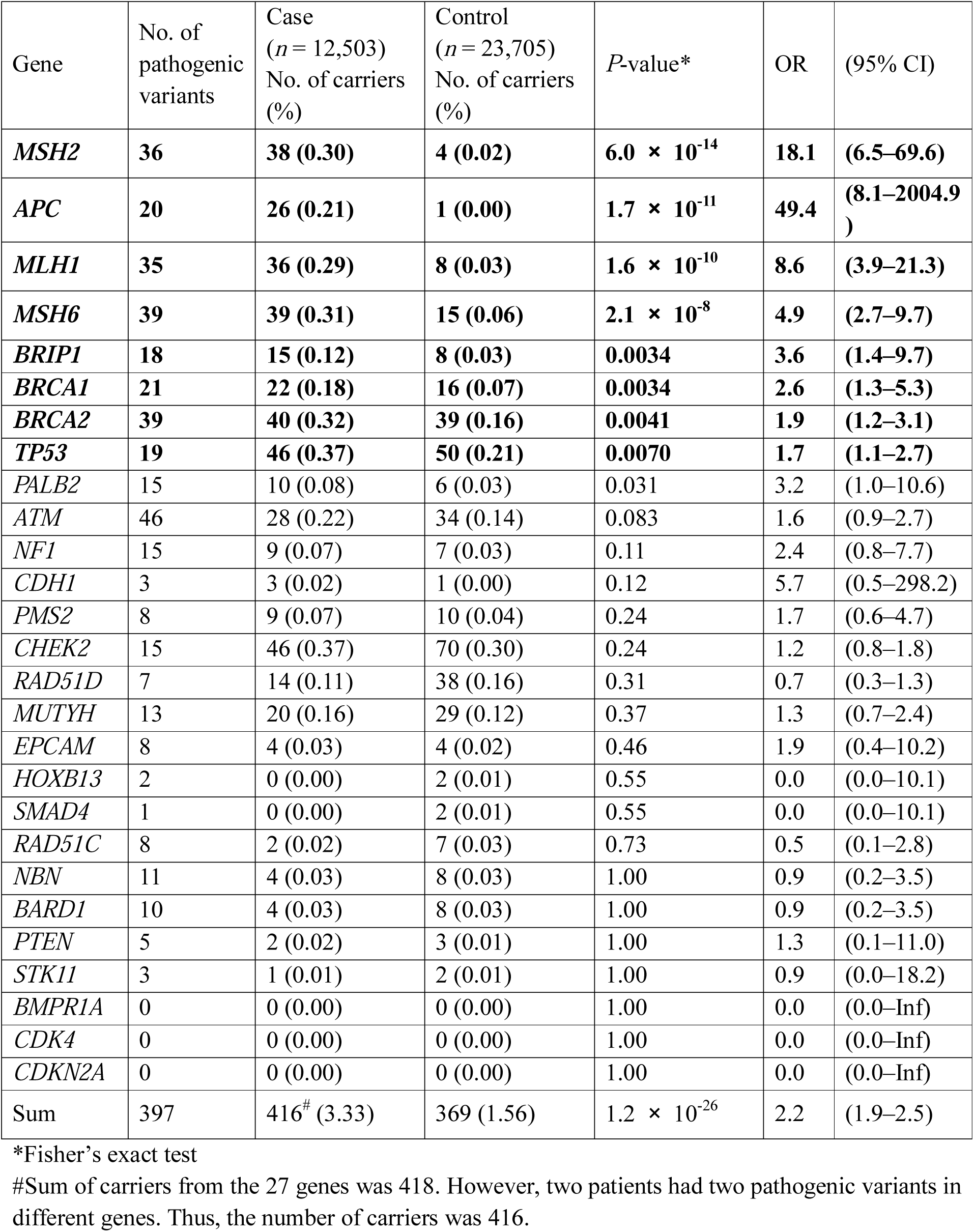
Result of gene-based association test using pathogenic variants.

### Statistical analysis

Case-control association analysis was performed using Fisher’s exact test under a dominant model. For *MUTYH*, we also used Fisher’s exact test under a recessive model. To investigate the association of pathogenic variants with demographic and clinical characteristics, we used *t*-tests for continuous variables and Fisher’s exact tests or Cochran-Armitage tests for discrete variables. All statistical tests were two-sided and *P* < 0.05 was considered statistically significant. For multiple hypothesis testing, *P*-values were adjusted using the Benjamini–Hochberg method, and a false discovery rate (FDR) < 0.05 was used as the significance threshold. All analyses were performed using the R statistical package (ver. 3.5.3).

### CNV analysis and MLPA

To screen putative pathogenic CNVs that affect MMR genes (*EPCAM, MSH2, MLH1, MSH6*) and *APC*, we performed CNV analysis with PennCNV (version 2013-02-08) (20) using Illumina high-density SNP microarray data for 12,246 and 23,392 controls previously obtained for a genome-wide association study (5). In our current cohort, four different SNP arrays were used for SNP genotyping (Human OmniExpress-24 v1.0, Human OmniExpressExome v1.0, v1.2, v1.4). Thus, we performed CNV calling by each array type separately. The signal intensity data of Log R Ratio and B allele frequency were exported from the Illumina BeadStudio software and were used as input for CNV calling. We used the default parameters and files, except for population frequency of B allele files, which we generated for each array type based on genotyping data of randomly selected Japanese subjects from BBJ (N = 1,000). We focused on deletions that overlapped or interrupted the coding sequences of target genes. For *EPCAM*, only deletions that affected exons 8 and 9 were recognized as pathogenic. For all putative pathogenic CNVs, we performed multiplex ligation-dependent probe amplification (MLPA) assays (21) using SALSA MLPA kits (Probe P003 and P043) on ABI3730xl.

### Quantitative PCR to evaluate the CNV of *EPCAM*

We performed qPCR to evaluate the CNV in exons 8 and 9 of *ECPAM* (22) in quadruplicate using an ABI7900HT instrument. The reaction mix included 10 ng of genomic DNA, TaqMan Gene Expression Master Mix, FAM-labeled probes (TGATGCATTTCAGTTATAC for exon 8; CACACACATTTGTAATTTG for exon 9), VIC-labeled probes for RNaseP, and primer pairs (ACTCCTAATCACTCTACCTTCCTACACA and CCTAAAGACAACAGTATAAAGGGACTC for exon 8; CTAACAAACTCATGACCTTCAAAGATG and AAAGGAGATGGGTGAGATGCA for exon 9). Thermocycling conditions were 50°C for 2 min and 95°C for 10 min, followed by 40 cycles at 95°C for 15 s and 60°C for 1 min. The SDS software (ABI) was used to evaluate the copy number of each exon.

## RESULTS

### Demographic and clinical characteristics of the subjects

After checking DNA quality and sequencing data, we analyzed a total of 12,503 CRC cases and 23,705 controls aged ≥ 60 years with no personal or family history of cancer from the disease-based biobank. The mean age at diagnosis was 65.1 years (SD 10.5) in the CRC cases. The percentage of males was 63.0% in the CRC cases and 52.6% in the controls, reflecting the fact that this proportion was the fifth smallest among the 42 male diseases registered in BioBank Japan (15), while the controls consisted of patients with complex diseases other than cancers in the biobank (13). A family history of CRC, gastric cancer, breast cancer, ovarian cancer, or prostate cancer was observed in 15.1%, 21.0%, 5.3%, 0.6%, and 2.4% of the CRC patient cohort, respectively. A personal history of gastric cancer, breast cancer, ovarian cancer, or prostate cancer was observed in 2.4%, 1.1%, 0.3%, and 1.1% of the CRC patient cohort. Other clinical characteristics are shown in **Table 1**.

### Pathogenic germline variants

Sequencing of the 27 hereditary cancer genes identified 4,804 germline variants in total. *BRCA2* and *APC* had the highest number of variants (548 and 495 variants, respectively) among the 27 genes (**Supplementary Figure 1a**). The region of *EPCAM-MSH2-MSH6* at chromosome *2p21* showed the highest number of variants per target sequence in the Japanese cohort (**Supplementary Figure 1b**). Among these variants, 2,096 were novel variants that were annotated in neither ClinVar nor dbSNP (build 150). We assigned their clinical significance using guidelines by the ACMG/AMP and ClinVar (criteria for classifying variants is **Supplementary Table 1 and 2**). Among them, 397 variants of the 27 genes were assessed as pathogenic, 941 as benign, and 3,466 as VUS (**Supplementary Figure 1c**). Among the 397 pathogenic variants, 199 (50.1%) were novel and not annotated in ClinVar.

**Figure 1.**
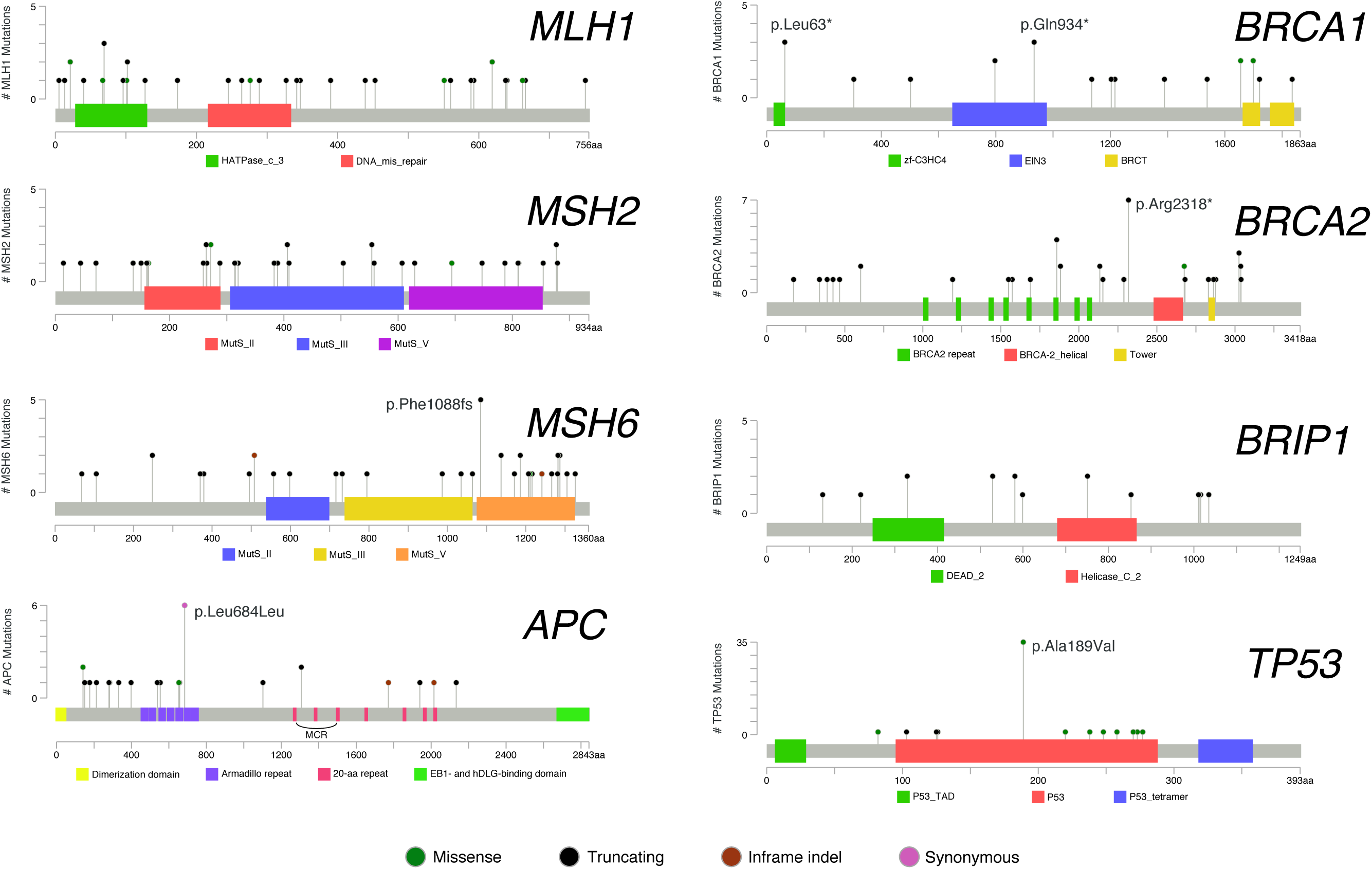
Location and the number of pathogenic variants in Japanese patients with CRC. Locations and protein domains of pathogenic variants found in CRC patients are shown by lollipop structures, with the variant type indicated by color. The x-axis reflects the number of amino acid residues, and the y-axis shows the total number of patients with each pathogenic variant. (a) *MLH1*, (b) *MSH2*, (c) *MSH6*, (d) *APC*, (e) *BRCA1*, (f) *BRCA2*, (g) *BRIP1*, and (h) *TP5*3. MutS_II: MutS domain II; MutS_III: MutS domain III; MutS_V: MutS domain V; MCR: mutation cluster region; HATPase_c_3: HSP90-like ATPase; DNA_mis_repair: DNA mismatch repair protein C-terminal domain; MutS_II: MutS domain II; MutS_III: MutS domain III; MutS_V: MutS domain V; zf-C3HC4: Zinc finger, C3HC4 type; EIN3: Ethylene insensitive 3; BRCT: BRCA1 C-Terminus domain; Helicase_C_2: Helicase C-terminal domain.P53_TAD: P53 transactivation motif; P53: P53 DNA-binding domain; P53_tetramer: P53 tetramerization motif.

We compared the frequency of carriers bearing pathogenic variants between CRC cases and controls (**Table 2**). In total, 3.3% of the CRC patients and 1.6% of the controls carried a pathogenic variant of the 27 genes (*P* = 1.20 × 10^−26^, odds ratio (OR) = 2.2). Pathogenic variants of eight genes (*MSH2, APC, MLH1, MSH6, BRIP1, BRCA1, BRCA2*, and *TP53*) were significantly enriched in the CRC cases (*P* < 0.01; FDR < 0.05), indicating their association with CRC development. We checked the locations and frequencies of the pathogenic variants (**Figure 1**) in the eight genes. Among the 227 variants in the eight genes, we observed four recurrent pathogenic variants shared in five or more Japanese CRC cases: *MSH6* p.Phe1088fs (n=5), *APC* p.Leu864Leu (n=6), *BRCA2* p.Arg2318* (n=7), and *TP53* p.Ala189Val (n=35). The prevalence of pathogenic variants of MMR genes was expected in the Japanese population with CRC, and the OR of pathogenic variants was 18.1 for *MSH2*, 8.6 for *MLH1*, and 4.9 for *MSH6*, which is consistent with the lower penetrance of CRC development in carriers with *MSH6* mutation (23, 24). Pathogenic variants of *MSH6* were clustered in the 3′ region, while no mutational clusters were observed in *MSH2* or *MLH1.* The OR of *APC* was 49.4, and pathogenic variants of *APC* in common CRC cases were clustered in its 5′ region, which suggests an attenuated phenotype of FAP (25, 26). Only one pathogenic variant, *APC* p.Glu1309fs, was detected in the mutation cluster region (MCR; 1,309–1,550 aa) of *APC* (25). This variant was found in two CRC patients, both of whom were diagnosed with CRC at a young age (27 and 31 years), although neither was formally diagnosed as FAP. In our cohort, two other CRC patients were diagnosed with FAP, which possessed pathogenic variants of *APC* outside the MCR (p.Arg283* and p.Ile1940fs).

Most interestingly, pathogenic variants of *BRIP1, BRCA1*, and *BRCA2* were significantly enriched in the CRC cases in the Japanese population. Several epidemiological studies have indicated the association of *BRCA1* and *BRCA2* with CRC development in the European population (27, 27, 29), but this association with CRC is still controversial and may be ethnicity-specific or variant-specific. *BRIP1* (*BRCA1-intereacting protein 1*) encodes a protein that is a member of the RecQ DEAH helicase family and interacts with BRCT repeat of BRCA1. This bound complex is essential for DNA double-strand break repair of the BRCA1 complex (30). We detected 15 CRC cases (0.12%) with pathogenic variants of *BRIP1* and 8 controls (0.03%) with pathogenic variants. Recurrent pathogenic variants of *BRIP1* have been reported to be associated with ovarian cancer (30) and breast cancer (31) in the European population, but its association with CRC in non-European populations is novel, although recurrent pathogenic variants of *BRIP1* are not found in the Japanese population. Four patients in our cohort had two pathogenic variants involved in BRCA-related genes. The clinical characteristics of these patients are shown in **Supplementary Table 3**.

**Table 3.** Demographic and clinical differences between CRC patients with and without pathogenic variants.

|  | No. of patients with pathogenic variants (%) | No. of patients without pathogenic variants (%) | *P*-value* | OR (95% CI) |
| --- | --- | --- | --- | --- |
| Number of subjects | 416 | 12,087 |  |  |
| Mean age at CRC diagnosis, y (SD) | 62.2 (12.0) | 65.2 (10.5) | $1.9 \times 10^{-6}$ | |
| Sex |  |  |  |  |
| Male | 275 (66.1) | 7,596 (62.8) | 0.18 | 1.2 (0.9–1.4) |
| Female | 141 (33.9) | 4,491 (37.2) |  | 1.0 (ref) |
| Personal history of gastric cancer | 16 (3.8) | 279 (2.3) | 0.048 | 1.7 (0.9–2.8) |
| Personal history of prostate cancer | 3 (0.7) | 129 (1.1) | 0.80 | 0.7 (0.1–2.0) |
| Personal history of breast cancer | 9 (2.2) | 129 (1.1) | 0.05 | 2.0 (0.9–4.1) |
| Personal history of endometrial cancer | 10 (2.4) | 36 (0.3) | $2.1 \times 10^{-6}$ | 8.2(3.6–17.1) |
| Personal history of ovarian cancer | 4 (1.0) | 37 (0.3) | 0.047 | 3.2 (0.8–8.9) |
| Personal history of FAP | 2 (0.5) | 0 (0.0) | 0.0011 | ∞ (5.5–∞) |
| Family history of CRC | 98 (23.6) | 1,791 (14.8) | $3.8 \times 10^{-6}$ | 1.8 (1.4–2.2) |
| Family history of gastric cancer | 101 (24.3) | 2,527 (20.9) | 0.099 | 1.2 (1.0–1.5) |
| Family history of prostate cancer | 13 (3.1) | 287 (2.4) | 0.33 | 1.3 (0.7–2.3) |
| Family history of breast cancer | 38 (9.1) | 622 (5.1) | 0.0011 | 1.9 (1.3–2.6) |
| Family history of endometrial cancer | 10 (2.4) | 112 (0.9) | 0.0075 | 2.6 (1.2–5.1) |
| Family history of ovarian cancer | 10 (2.4) | 59 (0.5) | $8.8 \times 10^{-5}$ | 5.0(2.3–10.0) |
| Family history of FAP | 1 (0.2) | 2 (0.0) | 0.10 | 14.5(0.2–279) |
| Anatomic site of CRC |  |  |  |  |
| Right-side colon | 129 (33.5) | 3,342 (29.7) | 0.087 | 1.2 (1.0–1.5) |
| Left-side colon and rectum | 248 (64.4) | 7,777 (69.0) |  | 1.0 (ref) |
| Both sides of colon | 8 (2.1) | 150 (1.3) | 0.16 | 1.7 (0.7–3.4) |
| Multiple CRC lesions |  |  |  |  |
| Yes | 36 (10.7) | 721 (7.4) | 0.035 | 1.5 (1.0–2.1) |
| No | 301 (89.3) | 8,989 (92.6) |  | 1.0 (ref) |
| Lymph node metastasis |  |  |  |  |
| Yes | 78 (40.4) | 2,504 (43.1) | 0.51 | 0.9 (0.7–1.2) |
| No | 115 (59.6) | 3,311 (56.9) |  | 1.0 (ref) |
| Distant metastasis |  |  |  |  |
| Yes | 11 (6.6) | 575 (12.5) | 0.022 | 0.5 (0.2–0.9) |
| No | 155 (93.4) | 4,031 (87.5) |  | 1.0 (ref) |

| Histology |  |  |  |  |
| --- | --- | --- | --- | --- |
| Adenocarcinoma |  |  |  |  |
| Papillary adenocarcinoma | 1 (0.3) | 63 (0.6) | 0.72 | 0.5 (0.0–2.8) |
| Tubular adenocarcinoma | 314 (86.7) | 9,358 (89.5) |  | 1.0 (ref) |
| Poorly differentiated adenocarcinoma | 18 (5.0) | 297 (2.8) | 0.024 | 1.8 (1.0–3.0) |
| Mucinous adenocarcinoma | 12 (3.3) | 150 (1.4) | 0.012 | 2.4 (1.2–4.3) |
| Signet-ring cell carcinoma | 0 (0.0) | 16 (0.2) | 1.0 | 0.0 (0.0–7.8) |
| Adenocarcinoma, NOS | 9 (2.5) | 386 (3.7) | 0.38 | 0.7 (0.3–1.4) |
| Adenosquamous carcinoma | 0 (0.0) | 19 (0.2) | 1.0 | 0.0 (0.0–6.4) |
| Squamous cell carcinoma | 1 (0.3) | 22 (0.2) | 0.53 | 1.4 (0.0–8.4) |
| Carcinoid tumor | 0 (0.0) | 43 (0.4) | 0.40 | 0.0 (0.0–2.7) |
| Others | 7 (1.9) | 106 (1.0) | 0.10 | 2.0 (0.8–4.2) |

\*Two-sided Fisher's exact test was used for all analyses except for age at diagnosis. A two-sided t-test was used for age at diagnosis. CI, confidence interval; OR, odds ratio; ref, reference group

### Clinical characteristics of patients with pathogenic variants

To investigate the association of pathogenic variants with the demographic and clinical characteristics of CRC, we compared the 416 carrier patients and the 12,087 non-carrier CRC patients (**Table 3**). The carriers were an average of 3.0 years younger at CRC diagnosis (62.2 years for carriers; 65.2 years for non-carriers; *P* = 1.9 × 10^−6^). Carriers more often had a family history of CRC (23.6% vs. 14.8%, *P* = 3.8 × 10^−6^), breast cancer (9.1% vs. 5.1%, *P* = 0.0011), endometrial cancer (2.4% vs. 0.9%, *P* = 0.0075), or ovarian cancer (2.4% vs. 0.5%, *P* = 8.8 × 10^−5^). Additionally, carriers tended to have a personal history of gastric cancer (3.8% vs. 2.3%, *P* = 0.048), endometrial cancer (2.4% vs. 0.3%, *P* = 2.1 × 10^−6^), or ovarian cancer (1.0% vs. 0.3%, *P* = 0.047). Carriers were also more likely to have multiple CRC lesions (10.7% vs. 7.4%, *P* = 0.035). Regarding CRC histological type, tumors of the carriers showed more mucinous adenocarcinoma (3.3% vs. 1.4%, *P* = 0.012) or poorly differentiated adenocarcinoma (5.0% vs. 2.8%, *P* = 0.024) than tumors of the non-carriers. We further examined the impact of pathogenic variants on age at CRC diagnosis (**Figure 2**). Pathogenic variants were found in 7.8% of patients diagnosed at <40 years of age. The proportion of pathogenic variants significantly decreased with advancing age at diagnosis (*P* = 1.2 × 10^−7^) but was stable between 2% and 3% in individuals 60 years of age or older.

**Figure 2.**
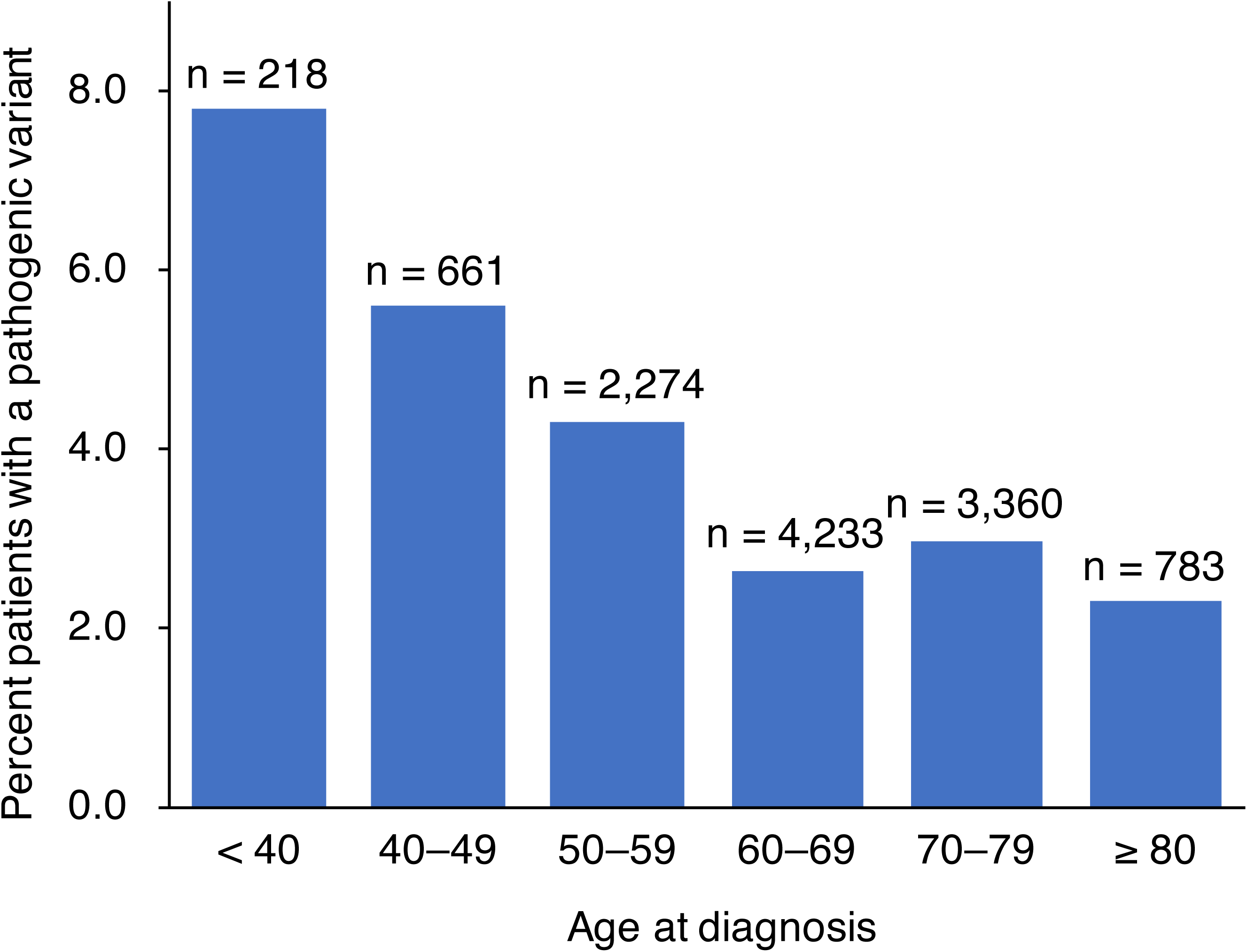
Proportion of patients with pathogenic variants by age at CRC diagnosis. The proportion of patients with pathogenic variants is shown. Two-sided Cochran-Armitage test was used (*P* = 1.2 × 10^−7^).

### Clinical characteristics of patients with functional categories of genes

We classified the 27 genes into five categories: MMR genes (*MLH1, MSH2, MSH6*, and *PMS2*), BRCA-related genes (*BRCA1, BRCA2*, and *BRIP1*), *APC, TP53*, and the remaining 18 genes. Pathogenic variants were found in the MMR genes of 122 patients, BRCA-related genes of 76 patients, *TP53* gene of 46 patients, *APC* gene of 26 patients, and other genes for 147 patients. Patients with pathogenic variants of MMR genes, *APC*, or *TP53*, were significantly younger at CRC diagnosis compared to patients without pathogenic variants (**Table 4**). Furthermore, pathogenic variants in each gene category were uniquely associated with clinical characteristics of patients (**Figure 3, Supplementary Table 4**). A family history of CRC was more frequent in patients with pathogenic variants of MMR genes (47.5%, *P* = 1.9 × 10^−17^) and *APC* (38.5%, *P* = 0.0028) in comparison to patients without pathogenic variants (14.8%). MMR genes were also associated with a family history of endometrial cancer (6.6% vs. 0.9%, *P* = 2.6 × 10^−5^) or ovarian cancer (3.3% vs. 0.5%, *P* = 0.0036), and personal history of endometrial cancer (8.2% vs. 0.3%, *P* = 2.1 × 10^−11^), which was mainly contributed by *MSH6*. CRC of MMR gene mutation carriers more often occurred in the right-side colon (49.1% vs. 30.1%, *P* = 2.8 × 10^−5^), had multiple lesions (16.0% vs. 7.4%, *P* = 0.0047), and was histologically classified as poorly differentiated adenocarcinoma (8.0% vs. 2.9%, *P* = 0.01) or mucinous adenocarcinoma (6.0% vs. 1.5%, *P* = 0.0043). In addition to family history of CRC, pathogenic variants of *APC* were associated with personal (7.7% vs. 0.0%, *P* = 4.4 × 10^−6^) and family (3.8% vs. 0.0%, *P* = 0.0064) history of FAP. Pathogenic variants of BRCA-related genes were associated with a personal history of breast cancer (6.6% vs. 1.1%, *P* = 0.0015) or ovarian cancer (3.9% vs. 0.3%, *P* = 0.002), and family history of breast cancer (19.7% vs. 5.1%, P = 7.5 × 10^−6^), but not CRC and prostate cancer. Pathogenic variants of the remaining gene group were significantly associated with a family history of ovarian cancer (P=0.0069, two variants of *CHEK2*, one of *ATM*, and one of *NF1)*.

**Table 4.**
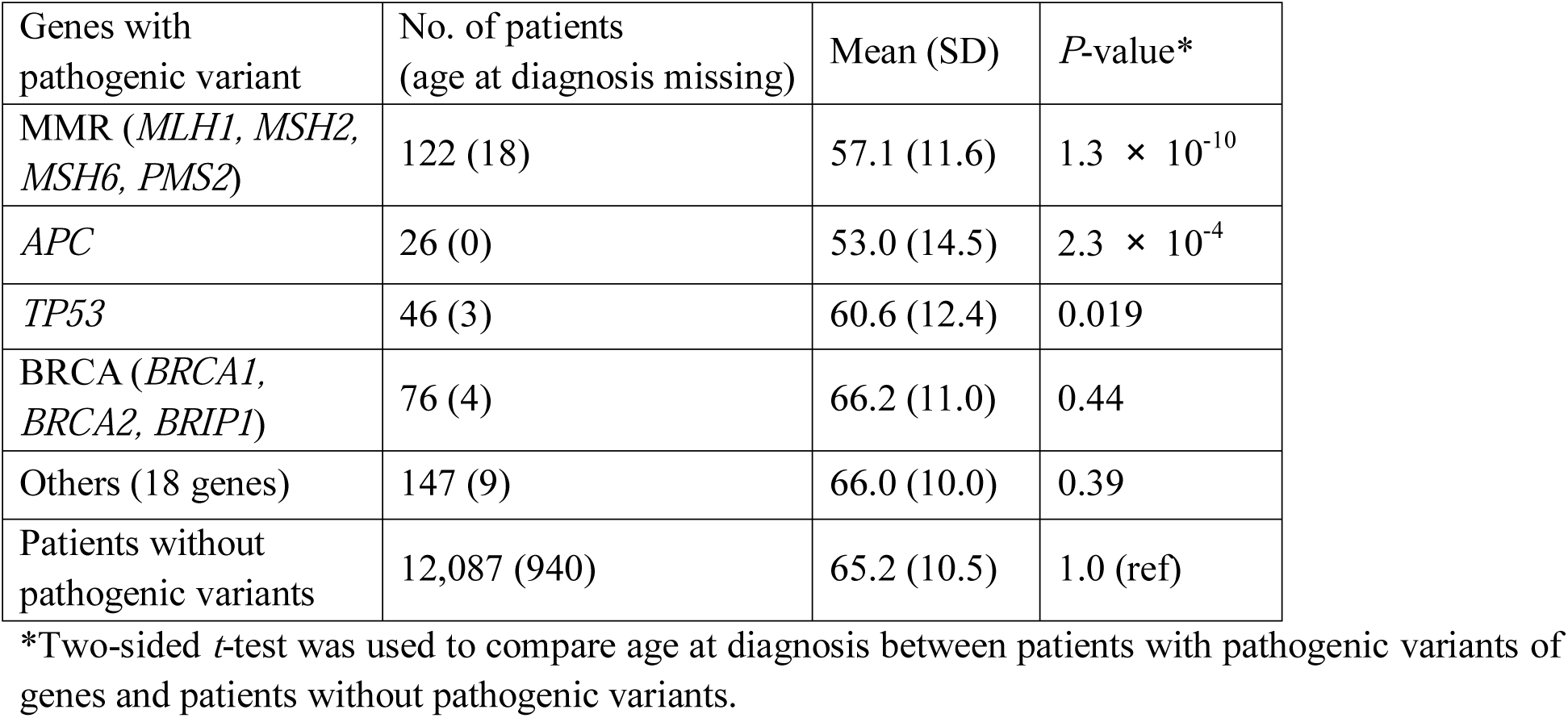
Mean age at CRC diagnosis in patients with pathogenic variants by gene category.

**Figure 3.**
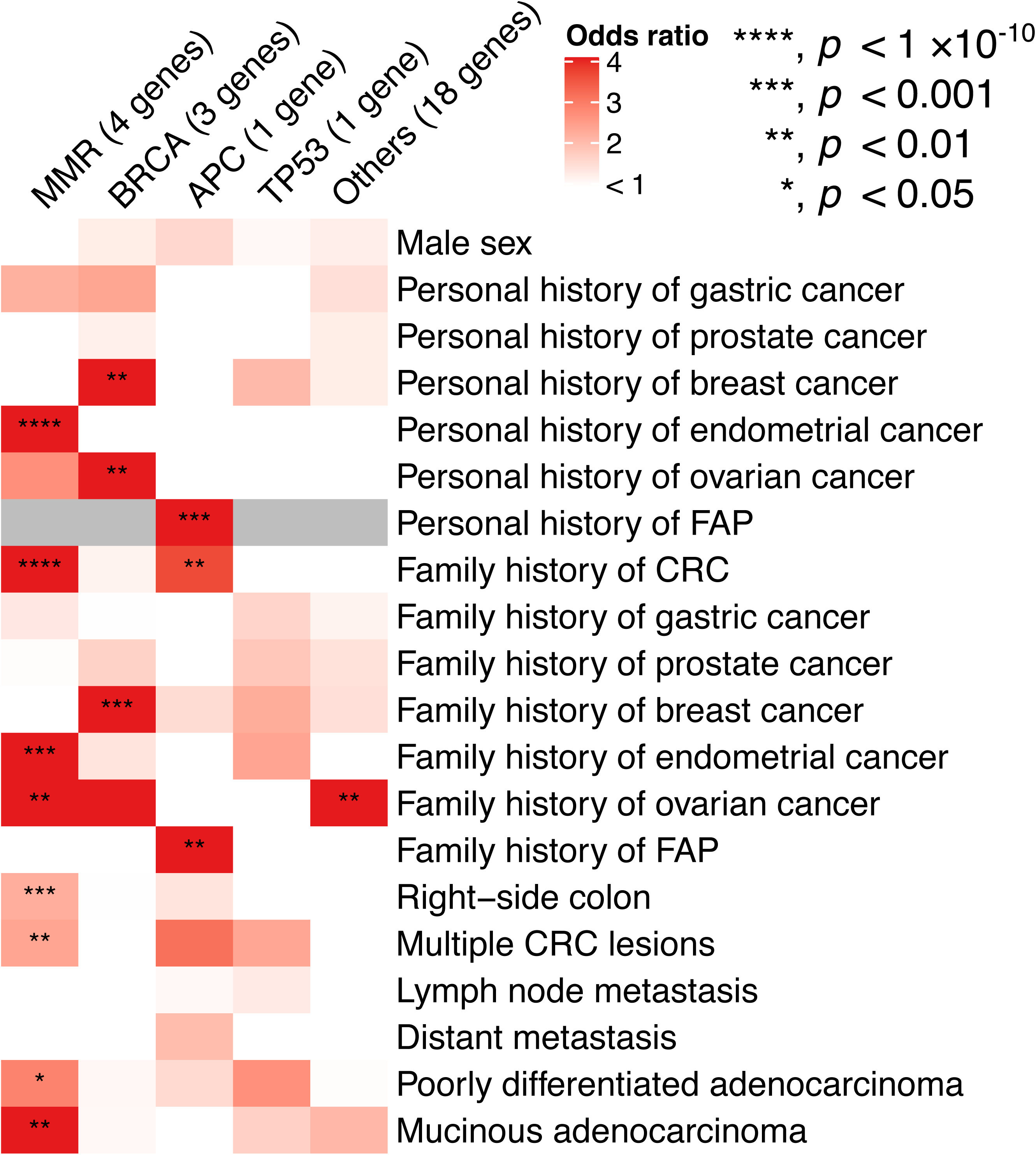
Statistical association between clinical characteristics and pathogenic variants by gene category. Odds ratios of clinical characteristics for patients with pathogenic variants are shown as a heatmap. Gray cells indicate an indefinite odds ratio. A two-sided Fisher’s exact test was applied using patients without pathogenic variants as the reference group. ****, *p* < 1 × 10^−10^; ***, *p* < 0.001; **, *p* < 0.01; *, *p* < 0.05.

### Clinical characteristics of patients with individual genes

When we examined the age at CRC diagnosis by gene, we found that the patients with pathogenic variants in *MSH2, MLH1, APC, TP53*, and *MSH6* were significantly younger at diagnosis compared to patients without pathogenic variants (**Supplementary Table 5**). In particular, patients with pathogenic variants in *MSH2, MLH1*, and *APC* were more than 10 years younger at diagnosis. The patients with pathogenic variants in *PALB2* were significantly older at diagnosis compared to patients without pathogenic variants.

### Copy number changes in MMR genes and *APC*

Copy number variants (CNVs) or structural variants (SVs) of *MSH2* and *MLH1* are responsible for 10–20% of Lynch syndrome cases (21, 32). In particular, the first exon of *MSH2*, last exon of *EPCAM*, and downstream of *EPCAM* are frequently affected by CNVs, likely due to the abundance of repetitive sequences around its gene body (33, 34). To identify CNVs of the hereditary cancer genes, we analyzed high-density SNP microarray data, which was previously obtained for GWAS of CRCs in 12,246 cases and 23,392 controls in BBJ (5). From this data, we extracted signal intensity and BAF information for the *MSH2* and *MSH6* regions at chromosome 2, *MLH1* at chromosome 3, and *APC* at chromosome 5, and called the CNVs of these genes in the CRC cases and the controls (98.56% and 98.60% were available). Among 35,600 samples, we detected 25 CNVs involving *EPCAM-MSH2, MLH1*, and *APC*, which were 16 and 2 deletions in the cases and controls, respectively, and 4 and 3 duplications, respectively (**Supplementary Table 6**). We also performed quantitative PCR targeting the last exon (exon 9) of *EPCAM* (32) and found five deletions among 12,502 CRC cases and 23,780 controls (**Supplementary Table 6**). Then, we performed MLPA assays on 24 DNA-available samples and validated 17 deletions and 6 duplications (12 for *EPCAM-MSH2*, 5 for *MLH1*, and 6 for *APC*), as shown in **Supplementary Figure 2**. In terms of clinical significance, large deletions of *MSH2, MLH1*, and *APC* would be pathogenic. Duplication involving a few exons, not including the last exon, are also likely pathogenic. But duplications of whole genes or large parts of the genes involving the last exon or the first exon are controversial and should be classified as VUS. We validated 23 CNVs of *MSH2, MLH1*, and *APC* in this cohort, 18 of which were assessed as pathogenic. We then combined pathogenic CNVs with pathogenic SNV/INDELs to assess their association between case and control. The combination of CNVs and SNV/INDELs decreased the OR compared to SNV/INDELs alone for *EPCAM-MSH2, MLH1*, and *APC* (**Supplementary Table 7**). However, statistical significance was improved for all genes.

In total, approximately 3.5% of CRC cases had a pathogenic variant or CNV of the 27 cancer-predisposing genes, while 1.3 % of the control population aged >60 years had a pathogenic variant or CNV in the Japanese population.

## Discussion

In this large-scale sequencing analysis targeting the 27 cancer predisposition genes, we found that pathogenic variants of *MLH2, MLH1, MSH6, APC, BRCA1, BRCA2, BRIP1*, and *TP53* were significantly enriched in CRC cases, indicating their association with CRC development. Pathogenic variants of *MLH2, MLH1*, and *MSH6* are associated with Lynch syndrome, and can lead to deficient DNA mismatch repair and development of MSI-high tumors in the colorectum, uterine, and other organs (6, 7). Among the four MMR genes, several studies demonstrated reduced penetrance for monoallelic carriers of *PMS2* variants compared with the other MMR genes (OR 2-5) (35) and our multiple-PCR based high-throughput sequencing method failed to cover some exons of *PMS2* because of the presence of several *PMS2* pseudogenes (36). Hence, we did not detect *PMS2* variants and our study may fail to obtain any evidence of *PMS2* association. Compared with *MSH2* and *MLH1, MSH6* showed a lower OR (OR=4.9), indicating a lower penetrance of *MSH6. MSH6* also showed a higher risk of endometrial cancer, as reported previously (23, 24). There are more ethnicity-specific variants in *MSH6* in the Japanese population (37). Here, we found recurrent variants of *MSH6* (p.Phe1088fs) in the 3′ region of *MSH6* (5 cases), and the pathogenic variants of *MSH6* tended to be enriched at the C-terminal region of MSH6, which plays a critical role in binding to MSH2 and the ATPase domain (38).

In this study, we identified many ethnicity-specific variants. For example, *MLH1* p.Leu582Val (45 cases and 102 controls in the Japanese population) and *MLH1* p.Thr151Thr (20 cases and 28 controls in the Japanese population) were annotated as pathogenic or VUS in ClinVar. However, since they were observed frequently in the Japanese non-cancer population, their annotations were changed to be benign. MSH6 p.Arg1076Cys is likely pathogenic in ClinVar, but it should also be benign because the frequency of this variant was found to be 0 in CRC cases and 4 in control cases in the Japanese population. This example demonstrates how variant information in non-European population facilitates annotation of rare variants. We should include more variant data in non-European populations to annotate hereditary cancer risk more precisely.

We detected many pathogenic variants of the *APC* gene in common CRC cases, most of which are truncated variants in the 5′ region of the *APC* gene, while two young-onset CRC cases had truncated variants in the MCR. Truncated variants in the 5′ region of *APC* genes are usually associated with attenuated FAP phenotype, and although the lifetime penetrance of CRC appears to be high in the attenuated FAP, CRC does not seem to develop in all affected patients (25, 26). In our study, some cases with pathogenic variants of *APC* were recorded to have few colorectal polyps with CRC, which was consistent with the concept of an attenuated phenotype. These pathogenic variants of *APC* are likely to retain some of the functions of *APC* protein by the mechanism of 5’ truncated mutation (39). The recurrent variants I1307K and E1317Q of *APC* confer increased risk of CRC in the Ashkenazi Jewish population (40), and we found one recurrent variant L684L (6 CRC cases and 0 control) of *APC* that was significantly enriched in Japanese CRC cases, although its biological significance is not clear. We did not detect any carriers with biallelic *MUTYH* pathogenic variants, whose mono-allelic mutation was also associated with CRC risk (41), and mono-allele pathogenic variants of *MUTYH* were not significantly associated with CRC in the Japanese population.

For *TP5*3, we detected 35 CRC cases carrying Ala189Val, which was reported in a case who developed multiple CRCs (42) and a case of late-onset Li-Fraumeni syndrome in the Asian population (43). Codon 189 resides in the L2 loop motif of the DNA-binding domain. *In silico* assessment of gene function predicted Ala189Val to be deleterious, but experimental data in yeast demonstrated that the Ala189Val mutant p53 retained fair trans-activity compared with wild-type *in vitro* (44). In this population study, the OR of *TP53* Ala189Val was relatively low (OR=1.7, case 35/12468 and control 39/23666), indicating its low penetrance for CRC development and more prevalence in Asian populations.

Most interestingly, we obtained evidence that pathogenic variants of *BRCA1, BRCA2*, and *BRIP1* are significantly enriched in CRC cases in the Japanese population. A recent meta-analysis of 14 studies (45) examining CRC risk in *BRCA1/2* carriers concluded that *BRCA1* mutation carriers had increased CRC risk (OR=1.49), whereas *BRCA2* mutation carriers did not (OR =1.09). *BRIP1* was reported to be associated with ovarian (30) and breast cancers (31), and one study suggested its association with CRC development in two cases (46). In our study, pathogenic variants of *BRIP1* showed a 3.6 OR, which is higher than *BRCA1/2*, indicating that it is worth being included on a list of panel sequencing to assess cancer risk, and a carrier with pathogenic variants of *BRIP1* should be subjected to CRC screening in addition to ovarian and breast cancers. Carriers with pathogenic variants of *BRCA1/2* and BRIP1 showed a greater family history of breast cancer (OR = 5.4) and ovarian cancer (OR = 6.3).

Copy number variants (CNVs) or structural variants (SVs) of *MSH2* and *MLH1* were responsible for 10-20% of Lynch syndrome (21, 32-34) and some FAP cases were also affected by CNV of APC (47). In addition to small-sized variants, we analyzed large-sized mutations, CNVs, by screening genome-wide SNP chip data and qPCR, and found 25 carriers with large deletions or duplications of *MSH2-EPCAM, MLH1*, and *APC*, but not *MSH6*. MLPA succeeded in validating 23 carriers, including 6 duplications. Large deletions involving exons or whole genes are obviously pathogenic, but duplications of whole genes or large parts of the genes are controversial (48). We should carefully evaluate the pathogenicity of these large duplications.

Regarding the phenotype of these pathogenic variants, we observed that onset age, personal cancer history, and family cancer history were apparently associated with carrier status of the pathogenic variants. *BRCA1/2* and *BRIP1* variants in CRC patients were associated with breast cancer and ovarian cancer, and MMR genes were associated with endometrial cancer. However, personal and family history of gastric cancer was not significantly associated with pathogenic variants, which is not consistent with the guideline that gastric cancer is on the tumor spectrum of Lynch syndrome. In Asian populations, gastric cancer is very prevalent due to infection by *Helicobacter pylori*, and its high incidence can affect the results of this population-based study in Japan.

In summary, we showed that the pathogenic variants in the Japanese CRC population were enriched in MMR genes, *APC, BRCA1/2, BRIP1*, and *TP53*, indicating that they can contribute to CRC development and risk. Universal screening of CRCs for various hereditary cancer syndromes, including Lynch syndrome, HBOC, and Li-Fraumeni syndrome, is feasible by NGS panel sequencing, and CRC risk should be assessed in multiple genes. This variant data in the Japanese population would contribute to the development of guidelines for genetic testing and variant interpretation for heritable CRCs. Variant data can also assist with universal screening, facilitate risk assessment, direct clinical management, and optimize screening strategies in Japanese and Asian populations.

## Acknowledgements

This research was supported by AMED under Grant Number JP19kk0305010. We thank the individuals who participated in this study. We acknowledge A. Sasaki, M. Mizukoshi, M. Endo, N. Hakozaki, A. Ogawa, and the staff of the BioBank Japan project and the RIKEN-IMS sequence platform for their technical assistances.

## Figure legends

**Supplementary Figure 1.**

(a) The number of variants in each of the 27 cancer-predisposing genes in this cohort. Orange, blue, and gray columns indicate pathogenic, benign, and VUS, respectively, as their clinical significance (determined using ACMG/AMP and pathogenicity assertions registered in ClinVar). (b) The number of variants per target sequence (1 kb) of each of the 27 cancer-predisposing genes in this cohort. (c) The ratio of pathogenic variants, benign variants, and VUS in each gene, which were annotated in this study.

**Supplementary Figure 1.**

MLPA validated large deletions and duplications of *EPCAM/MSH2* (a), *MLH1* (b), and *APC* (c). (a) *EPCAM*-exon 1-2 deletion, *EPCAM*-exon 1-5 deletion, *EPCAM* deletion, exon 7 deletion, exon 9-16 duplication, and whole duplication of *MSH2*. (b) Exon 12-13 deletion, exon 13-19 deletion, exon 1-5 deletion, and exon 3-10 duplication of *MLH1*. (c) Whole deletion, whole duplication, exon 1-16 duplication, and exon 17-18 duplication of *APC.*

## References

1. Jasperson KW, Tuohy TM, Neklason DW, and Burt RW. Hereditary and familial colon cancer. Gastroenterology 2010;138:2044–58.

2. Peters U, Bien S, Zubair N. Genetic architecture of colorectal cancer. Gut 2015;64:1623–36.

3. Jiao S, Peters U, Berndt S, et al. Estimating the heritability of colorectal cancer. Hum Mol Genet 2014;23:3898–905.

4. Graff RE, Möller S, Passarelli MN, et al. Familial risk and heritability of colorectal cancer in the Nordic twin study of cancer. Clin Gastroenterol Hepatol 2017;15:1256–64

5. Tanikawa C, Kamatani Y, Takahashi A, et al. GWAS identifies two novel colorectal cancer loci at 16q24.1 and 20q13.12. Carcinogenesis. 2018;39:652–60.

6. Lynch HT, and de la Chapelle A. Hereditary colorectal cancer. N Engl J Med. 2003;348:919–32.

7. Hampel H, Frankel WL, Martin E, et al. Screening for the Lynch syndrome (hereditary nonpolyposis colorectal cancer). N Engl J Med. 2005;352:1851–60.

8. Kinzler KW, and Vogelstein B. Lessons from hereditary colorectal cancer. Cell 1996;87:157–70.

9. Moreira L, Balaguer F, Lindor N, et al. Identification of Lynch syndrome among patients with colorectal cancer. JAMA.17;308:1555–65.

10. Hampel H, Pearlman R, Beightol M, et al. Assessment of tumor sequencing as a replacement for Lynch syndrome screening and current molecular tests for patients with colorectal cancer. JAMA Oncol. 2018;4:806–13.

11. Matthew B. Yurgelun, Matthew H. et al. Cancer susceptibility gene mutations in individuals with colorectal cancer. J Clin Oncol 35:1086–95, 2017

12. Jiang W, Cai MY, Li SY, et al. Universal screening for Lynch syndrome in a large consecutive cohort of Chinese colorectal cancer patients. High prevalence and unique molecular features. Int J Cancer 2019;144; 2161–68.

13. Hirata M, Kamatani Y, Nagai A, et al. Cross-sectional analysis of BioBank Japan clinical data: A large cohort of 200,000 patients with 47 common diseases. J Epidemiol 2017;27(3S):S9–S21.

14. Nagai A, Hirata M, Kamatani Y, et al. Overview of the BioBank Japan poject: Study design and profile. J Epidemiol 2017;27(3S):S2–S8.

15. Tamakoshi A, Nakamura K, Ukawa S, et al. Characteristics and prognosis of Japanese colorectal cancer patients: The Biobank Japan Project. J Epidemiol. 2017;27(3S):S36–S42.

16. Momozawa Y, Iwasaki Y, Parsons MT, et al. Germline pathogenic variants of 11 breast cancer genes in 7,051 Japanese patients and 11,241 controls. Nat Commun. 2018;9:4083.

17. Buys SS, Sandbach JF, Gammon A, et al. A study of over 35,000 women with breast cancer tested with a 25-gene panel of hereditary cancer genes. Cancer. 2017;123:1721–30.

18. Momozawa Y, Iwasaki Y, Hirata M, et al. Germline pathogenic variants in 7,636 Japanese patients with prostate cancer and 12,366 controls. J Natl Cancer Inst. 2019 Jun 19. pii: djz124.

19. Abou Tayoun AN, Pesaran T, DiStefano MT, et al. Recommendations for interpreting the loss of function PVS1 ACMG/AMP variant criterion. Hum Mutat. 2018;39:1517–24.

20. Wang K, Li M, Hadley D, et al. PennCNV: an integrated hidden Markov model designed for high-resolution copy number variation detection in whole-genome SNP genotyping data. Genome Res. 2007;17:1665–74.

21. Nakagawa H, Hampel H, and de la Chapelle A. Identification and characterization of genomic rearrangements of MSH2 and MLH1 in Lynch syndrome (HNPCC) by novel techniques. Hum Mutat. 2003;22:258.

22. Kuiper RP, Vissers LE, Venkatachalam R, et al. Recurrence and variability of germline EPCAM deletions in Lynch syndrome. Hum Mutat. 2011;32:407–14.

23. Baglietto L, Lindor NM, Dowty JG, et al: Risks of Lynch syndrome cancers for *MSH6* mutation carriers. J Natl Cancer Inst 2010;102:193–201.

24. Hendriks YMC, Wagner A, Morreau H, et al: Cancer risk in hereditary nonpolyposis colorectal cancer due to *MSH6* mutations: impact on counseling and surveillance. Gastroenterology 2004;127: 17–25.

25. Nieuwenhuis MH, and Vasen HF. Correlations between mutation site in APC and phenotype of familial adenomatous polyposis (FAP): a review of the literature. Crit Rev Oncol Hematol. 2007;61:153–61.

26. Soravia C, Berk T, Madlensky L, et al. Genotype-phenotype correlations in attenuated adenomatous polyposis coli. Am J Hum Genet. 1998;62:1290–301.

27. Grinshpun A, Halpern N, Granit RZ, et al. Phenotypic characteristics of colorectal cancer in BRCA1/2 mutation carriers. Eur J Hum Gent 2018;26:382–6.

28. Phelan CM, Iqbal J, Lynch HT, et al. Incidence of colorectal cancer in BRCA1 and BRCA2 mutation carriers: results from a follow-up study. Br J Cancer 2014;110:530–4.

29. Niell BL, Rennert G, Bonner JD, et al. BRCA1 and BRCA2 founder mutations and the risk of colorectal cancer. J Natl Cancer Inst. 2004;96:15–21.

30. Rafnar T, Gudbjartsson DF, Sulem P, et al. Mutations in *BRIP1* confer high risk of ovarian cancer. Nat Genet 2011;43:1104–07.

31. Seal S, Thompson D, Renwick A, et al. Truncating mutations in the Fanconi anemia J gene BRIP1 are low-penetrance breast cancer susceptibility alleles. Nat Genet. 2006;38:1239–41.

32. Wijnen J, van der Klift H, Vasen H, et al. MSH2 genomic deletions are a frequent cause of HNPCC. Nat Genet 1998;20:326–8.

33. Charbonnier F, Baert-Desurmont S, et al. The 5′ region of the MSH2 gene involved in hereditary non-polyposis colorectal cancer contains a high density of recombinogenic sequences. Hum Mutat 2005;26:255–61.

34. Ligtenberg MJ, Kuiper RP, Chan TL, et al. Heritable somatic methylation and inactivation of *MSH2* in families with Lynch syndrome due to deletion of the 3’ exons of *TACSTD1*. Nat Genet. 2009;41:112–7.

35. Senter L, Clendenning M, Sotamaa K, et al. The clinical phenotype of Lynch syndrome due to germ-line PMS2 mutations. Gastroenterology. 2008;135:419–28.

36. Nakagawa H, Lockman JC, Frankel WL, et al. Mismatch repair gene PMS2: disease-causing germline mutations are frequent in patients whose tumors stain negative for PMS2 protein, but paralogous genes obscure mutation detection and interpretation. Cancer Res. 2004;64:4721–7.

37. Terui H, Tachikawa T, Kakuta M, et al. Molecular and clinical characteristics of *MSH6* germline variants detected in colorectal cancer patients. Oncol Rep 2013; 30:2909–16.

38. Iaccarino, I., Marra, G., Palombo, F. & Jiricny, J. hMSH2 and hMSH6 play distinct roles in mismatch binding and contribute differently to the ATPase activity of hMutSalpha. EMBO J 1998;17, 2677–86.

39. Wein N, Vulin A, Falzarano MS, et al. Translation from a DMD exon5 IRES results in afunctional dystrophin isoform that attenuates dystrophinopathy in human and mice. Nat Med 2014;20:992–1000.

40. Frayling IM, Beck NE, Ilyas M, Dove-Edwin I, Goodman P, Pack K, Bell JA, Williams CB, Hodgson SV, Thomas HJ, Talbot IC, Bodmer WF, Tomlinson IP. The APC variants I1307K and E1317Q are associated with colorectal tumors, but not always with a family history. Proc Natl Acad Sci U S A. 1998;95:10722–7

41. Win AK, Dowty1 JG, Cleary SP, et al. Risk of colorectal cancer for carriers of mutations in *MUTYH*, with and without a family history of cancer. Gastroenterology. 2014;146:1208–1211.

42. Miyaki M, Iijima T, Ohue M, et al. A novel case with germline p53 gene mutation having concurrent multiple primary colon tumours. Gut. 2003;52:304–6.

43. Cho Y, Kim J, Kim Y, et al. A case of late-onset Li-Fraumeni-like syndrome with unilateral breast cancer. Ann Lab Med. 2013; 33:212–6.

44. Kato S, Han SY, Liu W, et al. Understanding the function-structure and function-mutation relationships of p53 tumor suppressor protein by high-resolution missense mutation analysis. Proc Natl Acad Sci U S A. 2003;100:8424–9.

45. Oh M, McBride A, Yun S, et al. BRCA1 and BRCA2 gene mutations and colorectal cancer risk: systematic review and meta-analysis. J Natl Cancer Inst. 2018;110:1178–89.

46. Ali M, Delozier CD, and Uzair Chaudhary U. BRIP-1 germline mutation and its role in colon cancer: presentation of two case reports and review of literature. BMC Med Genet. 2019; 20:75.

47. Kerr SE, Thomas CB, Thibodeau SN, et al. APC germline mutations in individuals being evaluated for familial Adenomatous polyposis: a review of the Mayo clinic experience with 1591 consecutive tests. J Mol Diag. 2013; 15:31–43.

48. Conner BR, Hernandez F, Souders B, et al. RNA analysis identifies pathogenic duplications in MSH2 in patients with Lynch syndrome. Gastroenterology 2019;156:1924–5.

