## Supplementary Figures for "Population-wide Screening for Germline Variants of Hereditary Cancer Genes in 12K Unselected Japanese Colorectal Cancers and 27K Controls"

### Slide 1
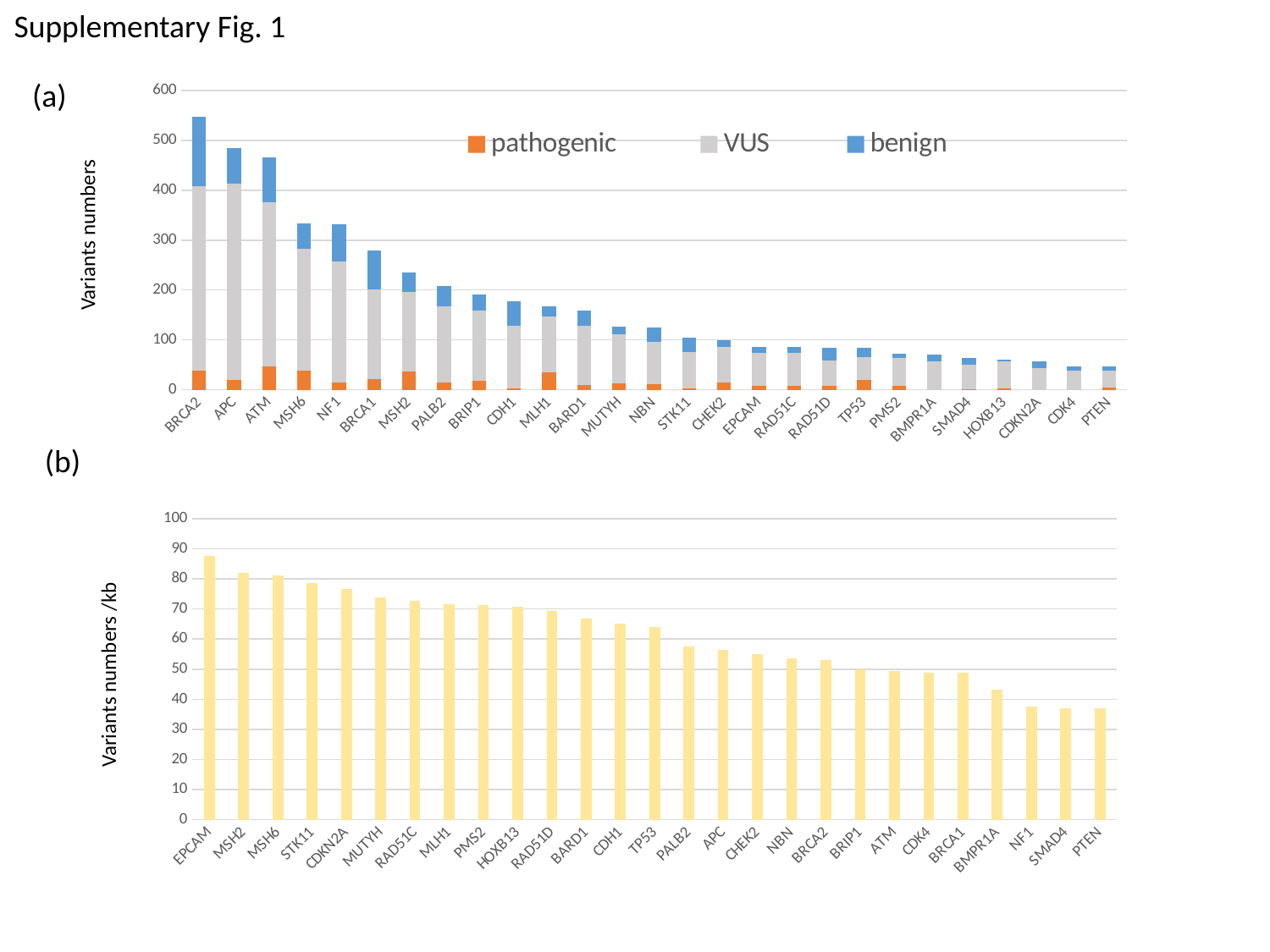

Supplementary Fig. 1
(a)
#### Chart
| Category | pathogenic | VUS | benign |
|---|---|---|---|
| BRCA2 | 39.0 | 369.0 | 140.0 |
| APC | 20.0 | 394.0 | 71.0 |
| ATM | 46.0 | 331.0 | 89.0 |
| MSH6 | 39.0 | 244.0 | 51.0 |
| NF1 | 15.0 | 242.0 | 75.0 |
| BRCA1 | 21.0 | 180.0 | 79.0 |
| MSH2 | 36.0 | 160.0 | 39.0 |
| PALB2 | 15.0 | 153.0 | 40.0 |
| BRIP1 | 18.0 | 141.0 | 32.0 |
| CDH1 | 3.0 | 126.0 | 48.0 |
| MLH1 | 35.0 | 112.0 | 21.0 |
| BARD1 | 10.0 | 118.0 | 31.0 |
| MUTYH | 13.0 | 98.0 | 15.0 |
| NBN | 11.0 | 85.0 | 29.0 |
| STK11 | 3.0 | 72.0 | 30.0 |
| CHEK2 | 15.0 | 70.0 | 15.0 |
| EPCAM | 8.0 | 66.0 | 12.0 |
| RAD51C | 8.0 | 66.0 | 11.0 |
| RAD51D | 7.0 | 52.0 | 25.0 |
| TP53 | 19.0 | 47.0 | 18.0 |
| PMS2 | 8.0 | 55.0 | 10.0 |
| BMPR1A | 0.0 | 56.0 | 15.0 |
| SMAD4 | 1.0 | 49.0 | 13.0 |
| HOXB13 | 2.0 | 54.0 | 5.0 |
| CDKN2A | 0.0 | 44.0 | 12.0 |
| CDK4 | 0.0 | 38.0 | 8.0 |
| PTEN | 5.0 | 34.0 | 7.0 |Variants numbers
(b)
#### Chart
| Category | |
|---|---|
| EPCAM | 87.66564729867483 |
| MSH2 | 81.91007319623563 |
| MSH6 | 81.00897404802328 |
| STK11 | 78.47533632286996 |
| CDKN2A | 76.60738714090287 |
| MUTYH | 73.90029325513196 |
| RAD51C | 72.5875320239112 |
| MLH1 | 71.58074137196421 |
| PMS2 | 71.2890625 |
| HOXB13 | 70.68366164542294 |
| RAD51D | 69.42148760330578 |
| BARD1 | 66.86291000841042 |
| CDH1 | 65.24143015112422 |
| TP53 | 64.07322654462243 |
| PALB2 | 57.569886520896766 |
| APC | 56.447858472998135 |
| CHEK2 | 54.91488193300385 |
| NBN | 53.671103477887506 |
| BRCA2 | 52.890647620886014 |
| BRIP1 | 49.92158912702561 |
| ATM | 49.47446650387514 |
| CDK4 | 48.93617021276596 |
| BRCA1 | 48.69565217391304 |
| BMPR1A | 43.21363359707851 |
| NF1 | 37.67162146828549 |
| SMAD4 | 36.9935408103347 |
| PTEN | 36.85897435897436 |Variants numbers /kb

### Slide 2
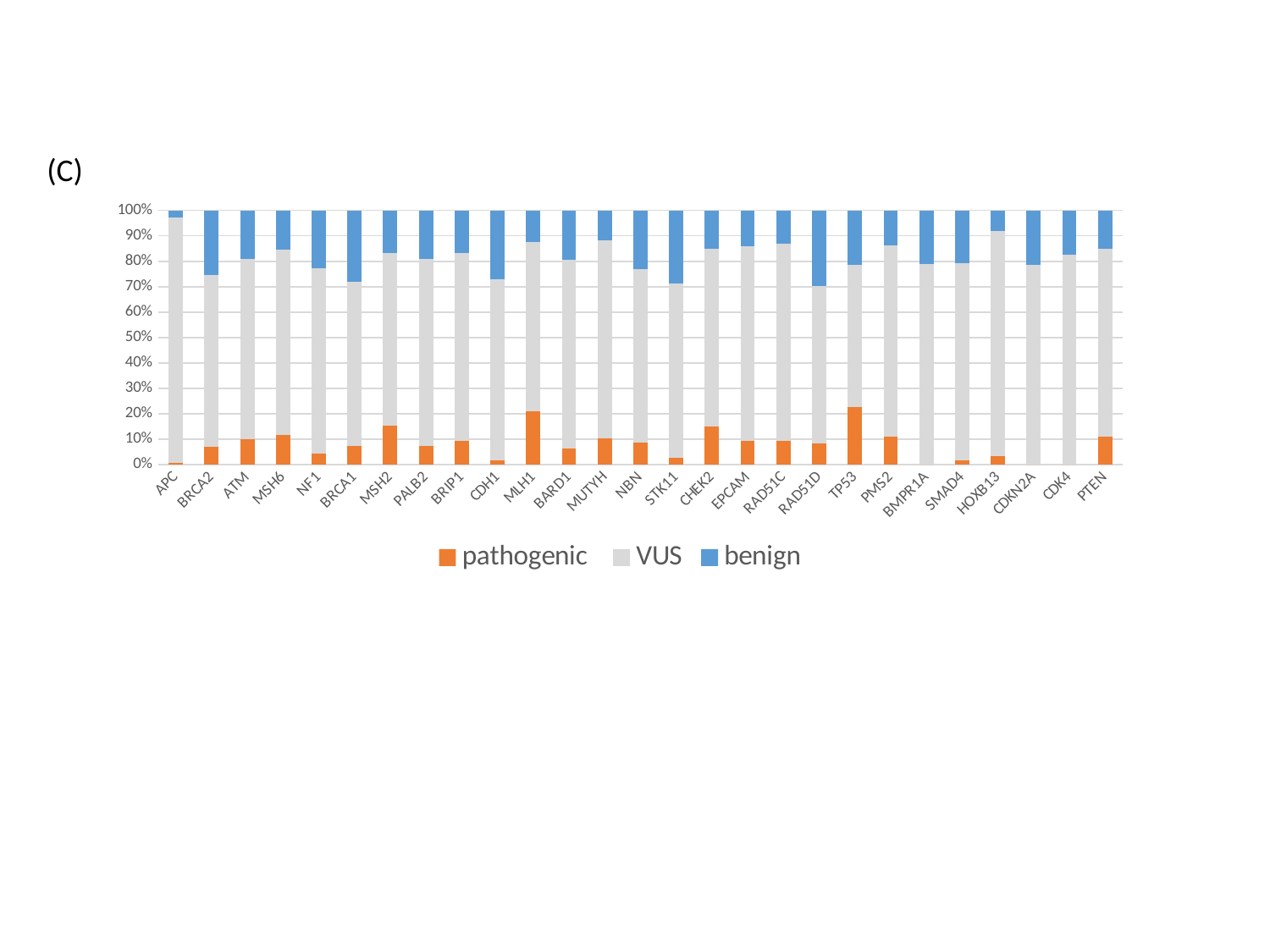

(C)
#### Chart
| Category | pathogenic | VUS | benign |
|---|---|---|---|
| APC | 20.0 | 2383.0 | 71.0 |
| BRCA2 | 39.0 | 369.0 | 140.0 |
| ATM | 46.0 | 331.0 | 89.0 |
| MSH6 | 39.0 | 244.0 | 51.0 |
| NF1 | 15.0 | 242.0 | 75.0 |
| BRCA1 | 21.0 | 180.0 | 79.0 |
| MSH2 | 36.0 | 160.0 | 39.0 |
| PALB2 | 15.0 | 153.0 | 40.0 |
| BRIP1 | 18.0 | 141.0 | 32.0 |
| CDH1 | 3.0 | 126.0 | 48.0 |
| MLH1 | 35.0 | 112.0 | 21.0 |
| BARD1 | 10.0 | 118.0 | 31.0 |
| MUTYH | 13.0 | 98.0 | 15.0 |
| NBN | 11.0 | 85.0 | 29.0 |
| STK11 | 3.0 | 72.0 | 30.0 |
| CHEK2 | 15.0 | 70.0 | 15.0 |
| EPCAM | 8.0 | 66.0 | 12.0 |
| RAD51C | 8.0 | 66.0 | 11.0 |
| RAD51D | 7.0 | 52.0 | 25.0 |
| TP53 | 19.0 | 47.0 | 18.0 |
| PMS2 | 8.0 | 55.0 | 10.0 |
| BMPR1A | 0.0 | 56.0 | 15.0 |
| SMAD4 | 1.0 | 49.0 | 13.0 |
| HOXB13 | 2.0 | 54.0 | 5.0 |
| CDKN2A | 0.0 | 44.0 | 12.0 |
| CDK4 | 0.0 | 38.0 | 8.0 |
| PTEN | 5.0 | 34.0 | 7.0 |

### Slide 3
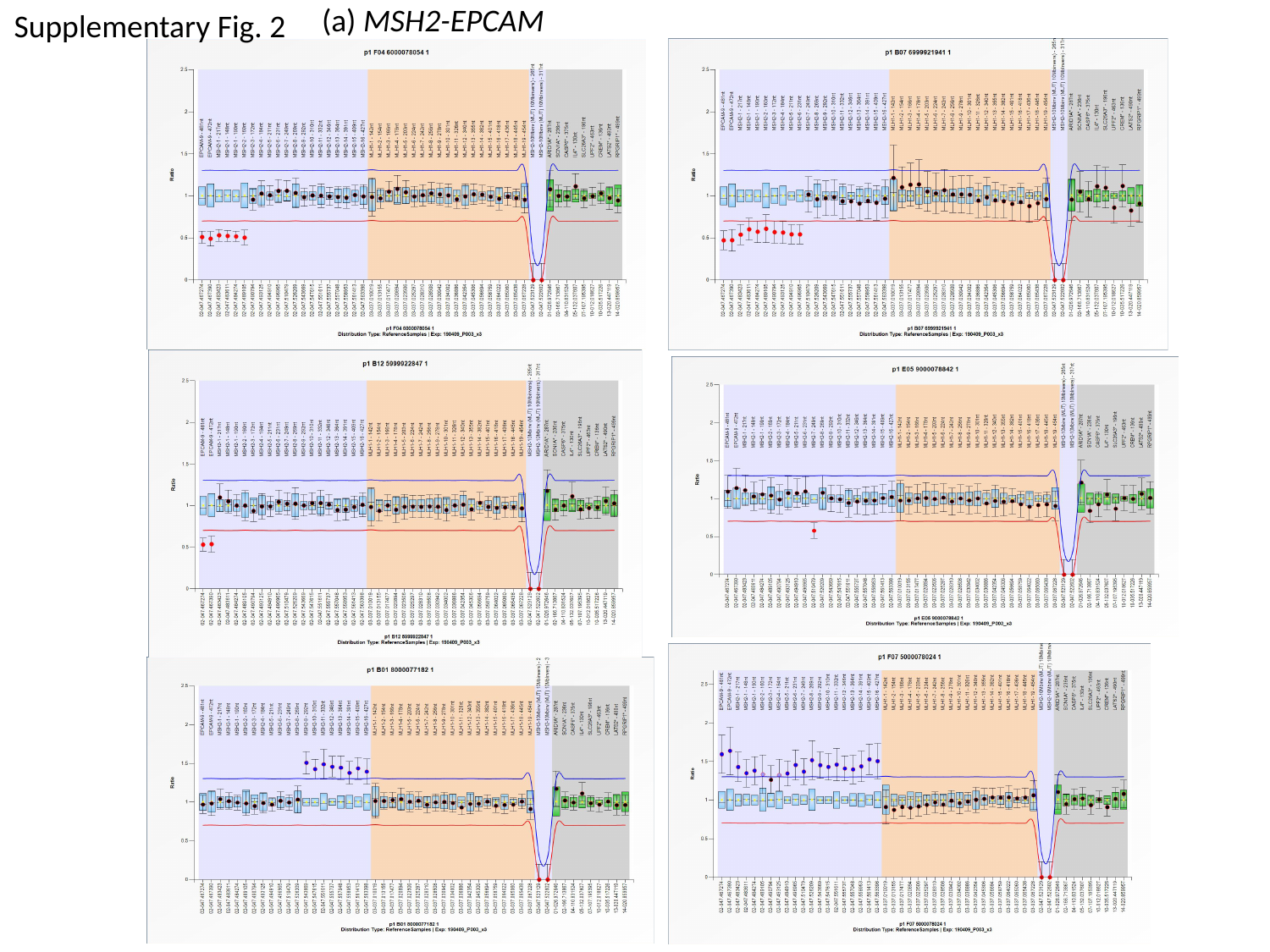

Supplementary Fig. 2
(a) MSH2-EPCAM

### Slide 4
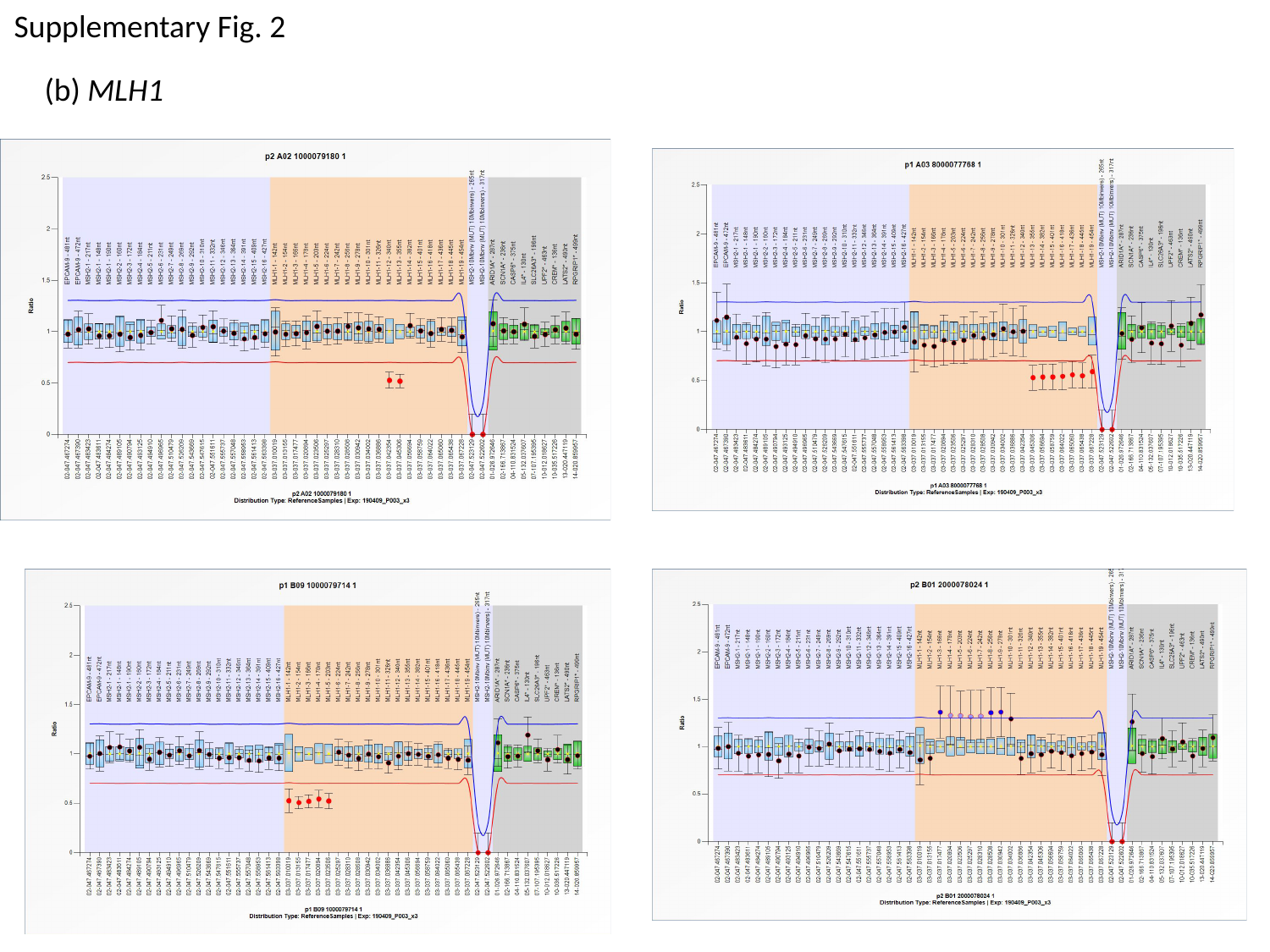

Supplementary Fig. 2
(b) MLH1

### Slide 5
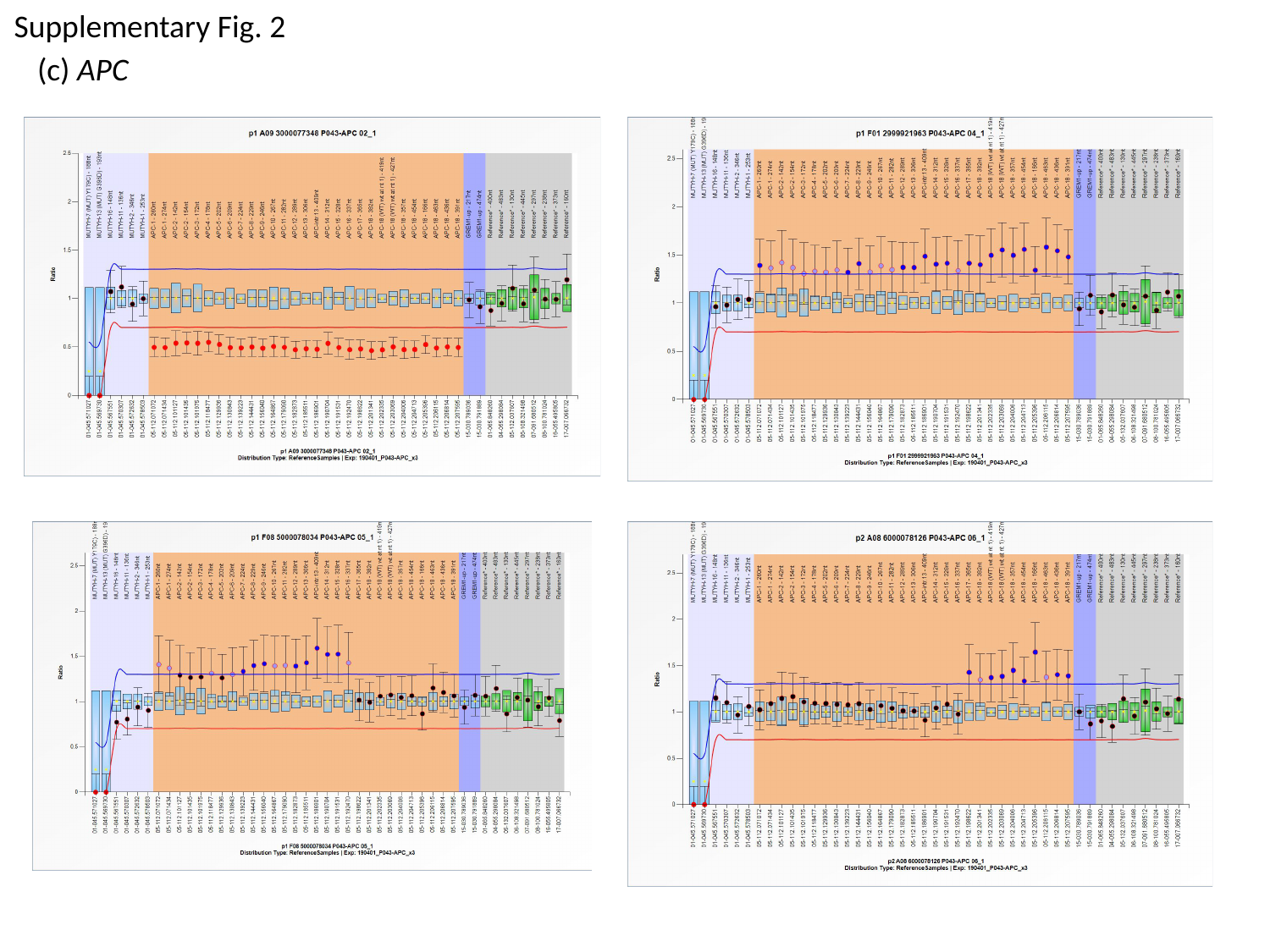

Supplementary Fig. 2
(c) APC
